# A retrospective molecular investigation of selected pigeon viruses between 2018-2021 in Turkey

**DOI:** 10.1101/2022.04.22.489161

**Authors:** Ismail Sahindokuyucu, Zafer Yazici, Gerald Barry

## Abstract

A recent first detection of pigeon aviadenovirus-1 and pigeon circovirus co-infection associated with Young Pigeon Disease Syndrome (YPDS) in a pigeon flock in Turkey, prompted a study focused on documenting the distribution of Pigeon aviadenovirus (PiAdV-1 and PiAdV-2), Pigeon circovirus (PiCV), Columbid alphaherpesvirus 1 (pigeon herpesvirus (PiHV)) and Fowl aviadenovirus (FAdV) in the country. These viruses were selected as they are associated with severe disease in pigeons across the world. A total of 192 cloacal swabs were collected from young (<1 year old) pigeons from 16 different private pigeon flocks across Turkey, between 2018 and 2021 as part of routine diagnostic sampling. PiCV genetic material was the most frequently detected 4/16 (25%), PiAdV-1 and CoHV-1 DNA were both found in one flock each, while neither PiAdV-2 and FAdV were detected in any of the studied pigeon flocks. PiCV and PiHV genetic material were both detected in the same pigeon flock’s cloacal samples as a co-infection with the identification of PiHV being a first in Turkey.

## Introduction

The common viral infections in young pigeons under 1 year old; are herpesviruses, adenoviruses and circoviruses. Viral infections of pigeons, particularly under the age of 1 year old, are associated with high morbidity and mortality. Young Pigeon Disease Syndrome (YPDS) a multifactorial disease, in which virus infections can play a role, is associated with high morbidity and mortality rates in the 3rd to 20th week of life. Clinically, signs tend to be quite non-specific with anorexia, depression, ruffled feathers, vomiting, diarrhoea, polyuria and a fluid filled crop being common [1].

Recently, a Pigeon aviadenovirus (PiAdV-1), Pigeon circovirus (PiCV) co-infection was documented in a Turkish pigeon flock, resulting in a loss of appetence, diarrhoea and high mortality rates [2]. It is possible that multiple infections in a flock could lead to increased disease burden in pigeons with increased potential for spillover infections. Pigeons are known transmitters of Newcastle Disease virus to chickens for example and spillover infections in wildlife have also been documented [3].

Columbid herpesvirus type 1, also known as pigeon herpesvirus (PiHV), is a member of the *Herpesviridae* family, and classified in the *Mardivirus* genus [4]. PiHV is regularly detected in pigeons worldwide, and is associated with depression, rhinitis, anorexia or conjunctivitis [5,6]. PiCV is a member of the genus *Circovirus* in the family *Circoviridae,* and pigeon infection was first documented in Canada in 1986 [7]. This virus is commonly associated with pigeons suffering from YPDS [1]. Pigeon aviadenovirus 1 and −2 (PiAdV-1 and −2) are members of the genus *Aviadenovirus* within the family *Adenoviridae* [8,9]. Fowl aviadenoviruses (FadV) have been previously isolated from both healthy and diseases pigeons [10–14]. It is not uncommon for viruses to be present and not cause disease. Bovine coronavirus, for example, can be present in the upper respiratory tract of healthy and diseased cattle. This does not mean the virus can’t be associated with disease but implies that other factors contribute to disease manifestation, be that co-infections with other pathogens or factors that induce stress in the animals [15,16]. Understanding the overall virome in a species enhances the ability to understand disease risk and manage it more effectively if it occurs, which is the motivation behind this study.

A select panel of viruses (PiCV, PiHV, PiAdV-1 and −2, and FadV) were chosen as they are the most common viruses of young pigeons, especially under the age of 1 year old and have been previously associated with YPDS [1,12,17]. Cloacal swabs taken from fancy pigeons were collected and tested for the presence of this panel of viruses, with confirmatory sequencing carried out to confirm the identifications.

## Materials and Methods

### Sampling

As part of routine diagnostics, to determine the cause of disease in flocks, 192 cloacal swabs from 16 different pigeon flocks were collected by sampling 12 randomly selected pigeons per flock (aged between 2 and 11 months of age) as part of routine diagnostics. The flocks were distributed across the Aegean part of Turkey and samples were collected between 2018 and 2021. All flocks had a history of dark green watery diarrhoea, inappetence and vomiting as well as sudden incidences of mortality (between 5 and 35% mortality). Samples were obtained from Izmir (10 flocks), Manisa (4 flocks) and Usak (2 flocks) provinces. Every flock was evaluated as one sample (12 cloacal swabs were pooled as one sample). Swabs were immersed in Dulbecco’s phosphate-buffered saline (DPBS, Thermo Fisher Scientifc), vortexed and stored at −20 °C until further analysis could take place.

### DNA Extraction and PCR

Swabs were defrosted on ice, trimmed using clean sterile scissors, vortexed again (1 minute) and centrifuged at 2000 x g for 10 min. The supernatants were then collected for screening by PCR.

Total DNA was extracted from 200 μl aliquots of prepared supernatants using a High Pure Viral Nucleic Acid Kit (Roche), according to the manufacturer’s protocol. The extracted samples were stored at −20 °C until PCR was performed.

The PCRs for PiAdV-1, PiAdV-2, PiHV, PiCV and FAdV were carried out in total volumes of 50 μl containing 10 μl of DNA, 0.8 μM of each primer, and PCR master mix (Xpert Fast Hotstart-Grisp, Portugal). The PCR for PiAdV-1, PiAdV-2 and FAdV was carried out in a thermocycler (Techne, UK) with an initial denaturation at 95°C for 5 min, followed by 35 cycles of 95°C for 20 seconds, 58°C for 30 seconds, and 72 °C for 1 min, with a final elongation step at 72 °C for 5 min. The PCR for PiCV and PiHV were carried out in the same thermocycler at the same time with an initial denaturation at 95°C for 5 min, followed by 35 cycles of 95°C for 20 seconds, 62°C for 30 seconds, and 72°C for 30 seconds, with a final elongation step at 72°C for 5 min. The PCR amplicons were visualised by agarose gel electrophoresis and then purified according to the manufacturer’s protocol (ExoSAP-IT PCR Product Cleanup Reagent, USA), before Sanger sequencing. All primers used in this study are listed in Table 1.

**Table 1.**
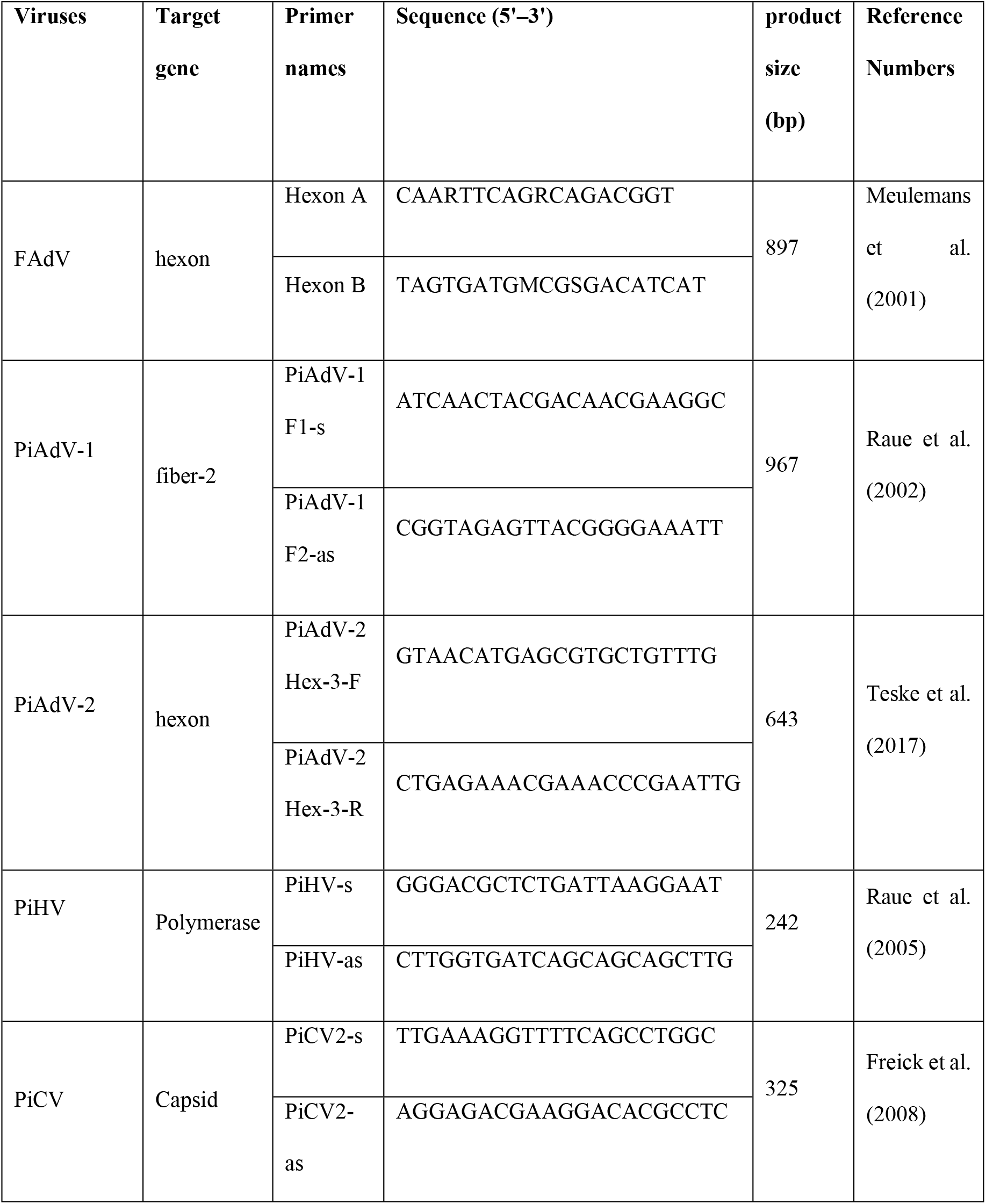
Information about the primers used in this study

Supernatants obtained from cloacal swabs were inoculated into the chorioallantoic cavity of 10-day-old specific pathogen-free (SPF) embryonated chicken eggs according to the protocol of the Villegas Laboratory Manual [18]. The eggs were incubated at 37 °C for 5 days and the allantoic fluid was then removed and inoculated into a fresh egg. This process was repeated until each supernatant was passaged through 5 eggs. After each passage an aliquot of the allantoic fluid was mixed with chicken red blood cells and incubated to test for the presence of viruses that cause haemagglutination.

### DNA sequencing and phylogenetic analysis

The PiAdV-1, PiCV and PiHV PCR amplicons were Sanger sequenced in both forward and reverse directions, using the same PCR primers as were used to amplify the products. Sequences, phylogenetic and molecular evolutionary analyses were conducted using MEGA version X (Molecular Evolutionary Genetics Analysis, version 10.1.8) software (The Biodesign Institute, Arizona, USA). The sequence data were then compared to most similar genome sequences and reference strains of aviadenovirus, herpesvirus and circovirus species from different countries which are available from the National Center for Biotechnology Information (NCBI) and their phylogenetic relationships were investigated. Phylogenetic trees were built using the neighbour-joining method with the T-92 (Tamura-3 parameter) [19] for pigeon pigeon aviadenoviruses and pigeon herpesvirus, K2 (Kimura-2 parameter with gamma distributed) (Kimura, 1980) for pigeon circovirus with substitution model using the maximum-likelihood statistical method in MEGA X, respectively [20].

#### Ethical statement

All samples were collected by veterinary professionals and analysed for diagnostic purposes, with the approval of the animal’s owners.

### Results

#### PCR screening results

In this study, a total of 16 pigeon flocks were analysed. The flocks were selected because of their widespread distribution and because breeders had reported that birds in each flock had clinical signs associated with YPDS and mortality in the flocks ranged from 5% to 35%. 4 out of 16 flocks (25%) tested positive for the presence of PiCV DNA, 1 out of 16 flocks (6.25%) tested positive for the presence of PiHV DNA and 1 out of 16 flocks (6.25%) tested positive for the presence of PiAdV-1 DNA. Of these positives, one flock had evidence of a co-infection of PiCV and PiHV as DNA fragments from both viruses were detected. FAdV and PiAdV-2 were not detected by PCR. Haemagglutination assays testing for the presence of viruses that could cause haemagglutination such as avian influenza or Newcastle Disease virus were negative.

The amplified fragments of the PiAdV-1 fiber-2 gene, the PiHV polymerase gene and the PiCV capsid gene were Sanger sequenced and BLASTed against the NCBI database (https://blast.ncbi.nlm.nih.gov) to compare these sequences with the genomes of aviadenovirus, herpesvirus and circovirus of pigeons. The PiCV fragments, now labelled, TR/PCV1, TR/PCV2, TR/PCV3 and TR/PCV4, matched most closely with isolates, KX108784 (China), MF664483 (Brazil), MT130538 (Turkey) and MK994767 (Poland) respectively (Fig 1).

**Fig 1.**
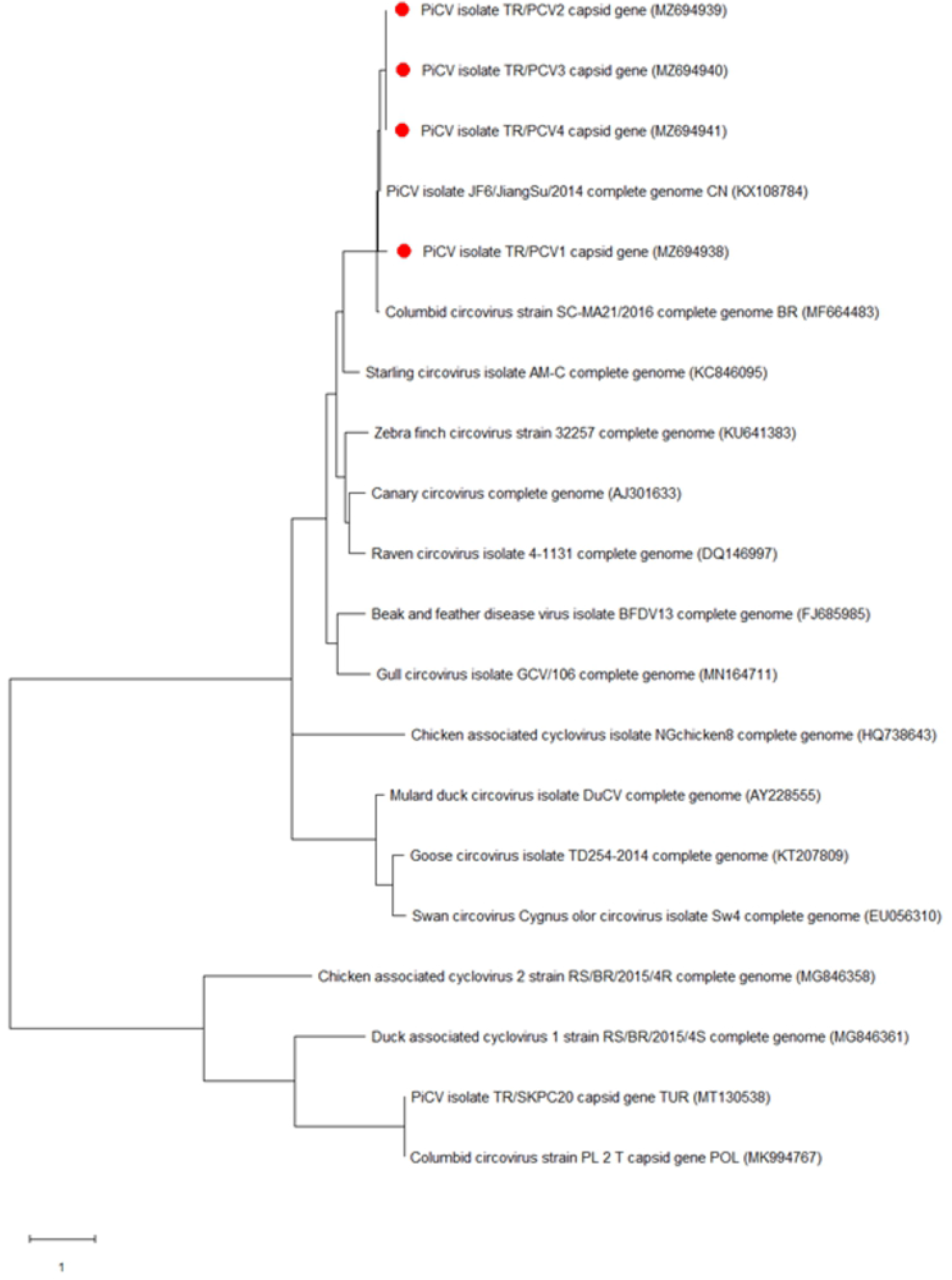
Phylogenetic tree based on the nucleotide sequence of the PiCV capsid gene. The tree shows circovirus species which affect fowl. The tree shows the names and GenBank accession numbers of each isolate. The PiCV isolates from this study are indicated by a red dot.

The PiAdV-1 sequence, now designated TR/PADV1, was found to be closely related to a previously sequenced PiAdV-A strain called TR/SKPA20 (Turkey, 99.46% similar) and another called P18-05523-66 (Australia, 98.82% similar) and while it was 98.5% similar to IDA4 (Netherlands) at the amino acid level (Fig 2).

**Fig 2.**
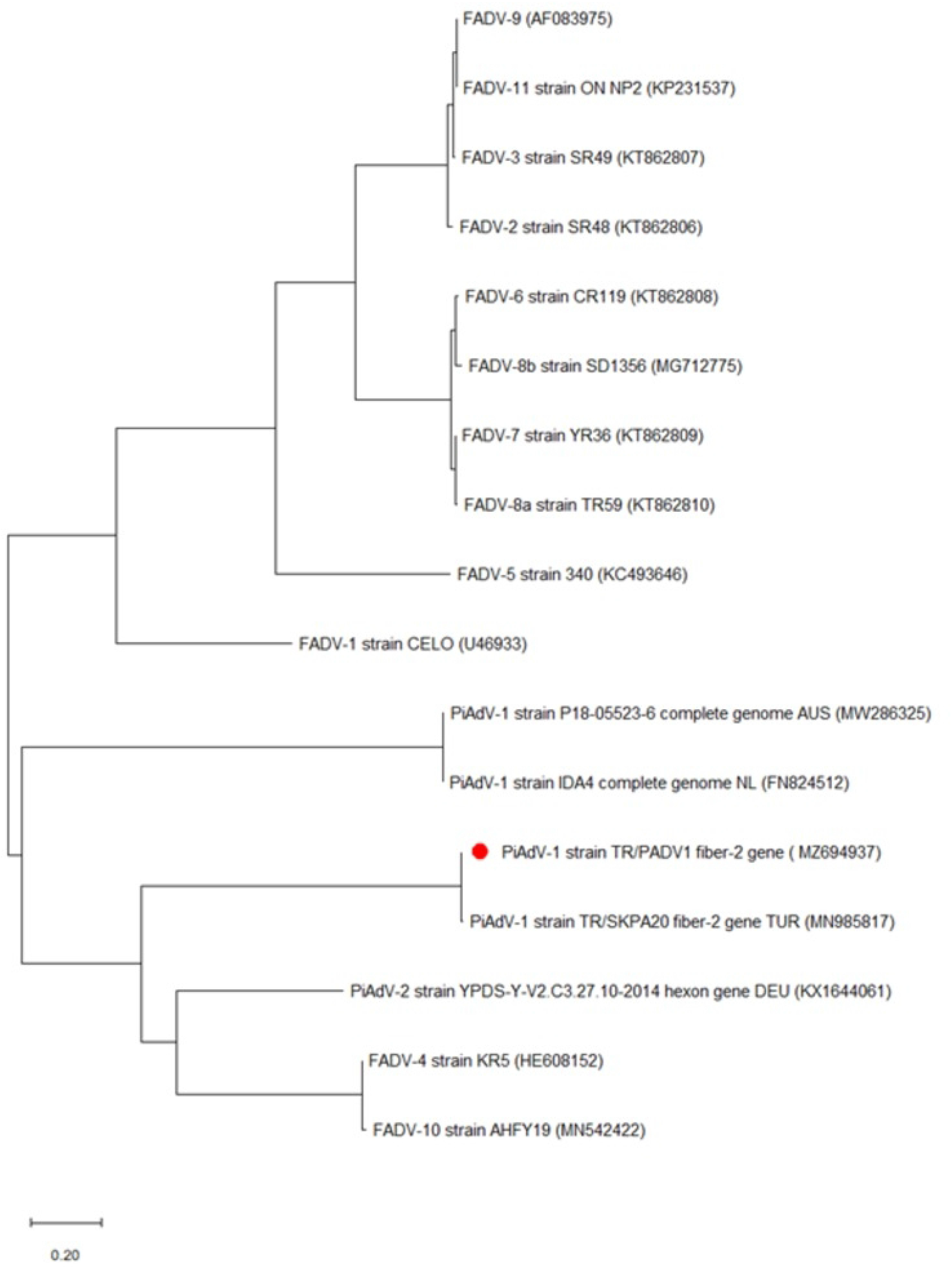
Phylogenetic tree based on the nucleotide sequence of the PiAdV-1 fiber-2 gene. The tree shows adenovirus species which affect fowl. The tree shows the names and GenBank accession numbers of each isolate. The PiAdV isolate from this study is indicated by a red dot.

The PiHV sequence, now designated TR/PHV1, was found to be most closely related to strains previously isolated in China (KX589235) with 98.77% and Thailand (KM010015) with 98.76% similarity at the amino acid level (Fig 3). The nucleotide sequences of the field isolates presented here (TR/PCV1, TR/PCV2, TR/PCV3, TR/PCV4, TR/PADV1 and TR/PHV1) have been submitted to the NCBI-GenBank database under the following accession numbers: (MZ694938, MZ694939, MZ694940, MZ694941, MZ694937 and MZ694942).

**Fig 3.**
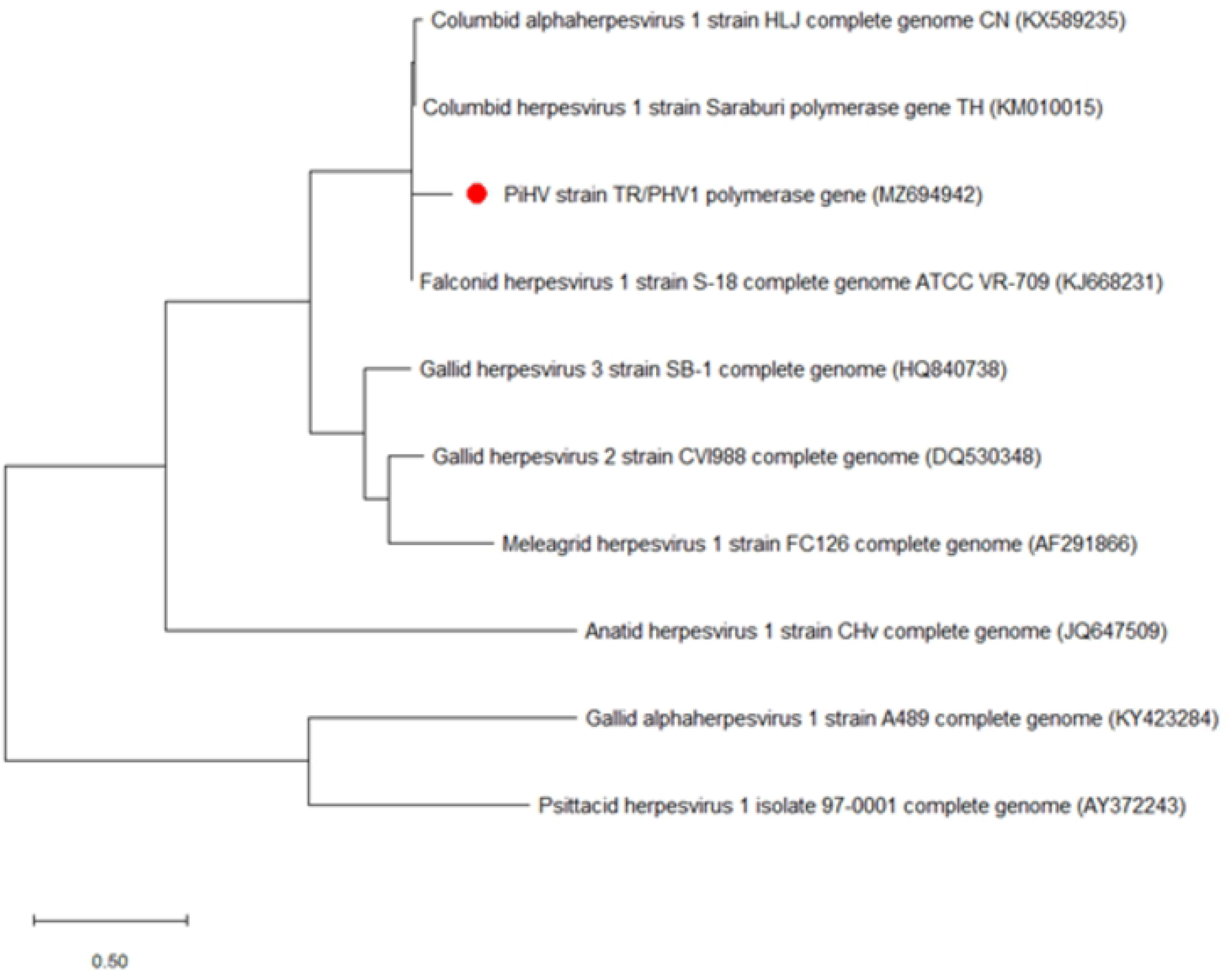
Phylogenetic tree based on the nucleotide sequence of the PiHV polymerase gene. The tree shows herpesvirus species which affect fowl. The tree shows the names and GenBank accession numbers of each isolate. The PiHV isolate from this study is indicated by a red dot.

## Discussion

This study describes a molecular retrospective survey for the detection of selected pigeon DNA viruses; PiHV, FAdV, PiCV and PiAdV. These viruses were selected as they are the most important viruses of young pigeons and have been regularly associated with YPDS, a major disease in young pigeons [1,12,17].

PiHV, FAdV, PiCV and PiAdV infections are clinically indistinguishable in pigeons, and reliable diagnostic tests are needed to identify these viruses correctly. PCR is the most common diagnostic test for each; PiHV by targeting regions of the DNA polymerase gene [1,4,21], the 12 serotypes of FAdV by amplifying a conserved pedestral region of the hexon gene [22,23], PiCV by detecting the capsid gene [1,24] or highly conserved genomic regions [25], and PiAdV by detecting the fiber-2 and hexon gene [9,26]. In this study, a combination of PCR and Sanger sequencing was used to confirm each positive result.

Adenoviruses have been previously documented to infect pigeons with younger birds less than 1 year old appearing to be particularly vulnerable to disease [27]. FAdV serotypes, for example, have also been identified from diseased pigeons but interestingly have also been found in healthy pigeons, indicating that other factors may also contribute to disease manifestation [10,12,28]. PiAdV-1 although similar to FAdV, has been identified, and distinguished from the 12 FAdV serotypes by cross-neutralization assays [28]. These tests were subsequently supported by genome sequencing, confirming PiAdV-1 as a new species in the genus Aviadenovirus [29]. It is not surprising to identify it among the flocks that were tested, although the prevalence (1 out of 16 flocks) suggests that it is not creating a major burden of infection in the fancy pigeon populations of Turkey.

In general, circovirus infections in birds are associated with immunosuppression, feather disorders and low weight. Immunosuppressive viruses tend to exacerbate the impact of other infections and can be major contributors to polymicrobial associated conditions such as YPDS. PiCV is believed to cause immunosuppression in pigeons through increased apoptosis in B lymphocytes and this is likely to facilitate secondary infections or amplification of pathogens that are already present [30]. Viral and bacterial co-infections may aggravate the clinical picture of other infections especially in pigeon lofts. This situation has also been shown for other avian and mammalian species with Bovine respiratory disease being a perfect example where multiple bacteria and viruses are associated, and likely exacerbate each others impact [31,32]. A number of PiCV prevalence studies have been carried out in different parts of the world although none to date in Turkey. Initially, prevalence studies based on histological analysis of tissue samples from diseased birds revealed relatively low (less than 50%) prevalences in populations but with increased use of molecular methods with increased sensitivity the virus was regularly found in almost all birds that were tested and was also commonly found in apparently health birds. Diseased young birds less than 1 year old were most likely to be infected, with one study in Poland showing 100% of birds tested to be infected with PiCV [33]. Work in China similarly identified PiCV DNA in 80.7% of diseased birds that were tested and 63.4% of asymptomatic birds that were tested [34]. In this study, the prevalence of PiCV (25%) among clinically affected pigeon flocks is lower than in these and many other studies. It is unclear why this might be the case, but, because genetically the birds in Turkey are similar to birds globally, there is unlikely to be an inherent resistance to this virus and therefore suggests PiCV infection is likely to increase over time and may lead to an increase in the burden of disease in the future [35–38].

PiHV is frequently found in raptors as well as pigeons. A study in the UK suggested that PiHV prevalence ranges from 4.9% within the *Strigidae* to 8.8% in the *Accipititridae* family [39]. Seroprevalence data from the same report identified antibodies against PiHV in 8.2% of free-ranging owls. This suggests that this herpesvirus has the ability to infect a number of different species of bird and therefore is likely to be able to move between species quite readily. In this study, according to the BLAST analysis of the polymerase gene of TR/PHV1, the virus material identified is very similar to that previously found in raptors suggesting that cross species transmission is happening in Turkey. This is the first identification of this virus in Turkey and transmission from migratory birds can not be ruled out as a possible source. This may occur through close physical contact or potentially through shared food sources or habitat linked fomite transmission. This should be a biosecurity consideration for owners of pigeons going forward.

In this study, we have identified the presence of PiAdV-1, PiCV and PiHV DNA in pigeon flocks in Turkey, including one flock where both PiCV and PiHV DNA were found suggestive of a co-infection. These findings act as a first prevalence study in Turkey for these viruses and suggest that they are contributing to the infectious burden of pigeons and possibly other birds in this region. This study contributes to increased knowledge of the virome of pigeons in Turkey and should inform the biosecurity approaches of owners.

## Acknowledgements

We thank Mr. Bahtiyar Yilmaz from Letgen, Biotechnology for technical support.

## Author contributions

IS collected and analysed the samples. IS, ZY and GB analysed the data generated. IS, ZY and GB wrote the manuscript.

## Declarations

### Availability of data and materials

Sequences from the study have been deposited in the GenBank database and are freely accessible.

